# A neuronal population in the bed nucleus of the stria terminalis mediating pup-directed aggression in male mice

**DOI:** 10.64898/2026.02.26.708360

**Authors:** Kengo Inada, Haruaki Sato, Mitsue Hagihara, Kazunari Miyamichi

## Abstract

The mammalian brain orchestrates a broad range of social behaviors, including interactions with conspecific young. Sexually inexperienced male mice typically exhibit pup-directed aggression, whereas fathers display caregiving behaviors. Although this behavioral transition provides a valuable framework for understanding life stage-dependent modulation of social behavior, the underlying mechanisms remain unclear. The posterior bed nucleus of the stria terminalis (BST), particularly its principal subdivision (BSTpr), is a critical node regulating pup-directed aggression. Within this region, neurons expressing *estrogen receptor 1* (*Esr1*) promote pup-directed aggression in female mice; however, the specific neuronal subtypes mediating this behavior in males remain unknown. Here, reanalysis of single-cell transcriptomic datasets shows that neurons expressing (+) *cocaine- and amphetamine-regulated transcript prepropeptide* (*Cartpt*) constitute a subset of *Esr1*+ neurons within the BST. Exposure to pups selectively increased *c-fos* expression, a marker of neuronal activation, in BSTpr *Cartpt*+ neurons in infanticidal virgin males. Chemogenetic activation of these neurons enhanced pup-directed aggression, whereas their ablation suppressed infanticide. Anatomical analyses demonstrated that BSTpr *Cartpt*+ neurons received direct inhibitory inputs from the anterior commissure nucleus (ACN), a region associated with parental behavior. Additionally, inhibitory transmission from the ACN to BSTpr *Cartpt*+ neurons was enhanced in fathers. Together, these findings identify BSTpr *Cartpt*+ neurons as key mediators of pup-directed aggression in male mice, suggesting a neural circuit mechanism underlying the transition from aggression to parental care.

## Introduction

The mammalian brain orchestrates a wide range of social behaviors, including interactions with conspecific young. Notably, the degree of aggression toward offspring changes depending on the life stage and experience. For example, virgin male mice typically attack unfamiliar pups, whereas fathers display caregiving behaviors (1-4). Understanding the mechanisms underlying this behavioral transition is essential for elucidating the neural control of social behavior; however, these mechanisms remain unclear.

Over the past decade, studies have identified specific limbic neurons and circuits governing either infanticidal or paternal caregiving behaviors in male mice. Neurons in the medial preoptic area (MPOA) expressing *galanin, Esr1*, or *calcitonin receptor* function as parental behavior centers (5-8). In contrast, GABAergic neurons in the posteroventral subdivision of the medial amygdala, *urocortin-3*-expressing (+) neurons in the perifornical area, and excitatory projection neurons from the amygdalohippocampal area to the MPOA are associated with infanticide (9-11). Furthermore, the posterior bed nucleus of the stria terminalis (BST), including regions such as the BSTpr and the rhomboid nucleus, plays a role in pup-directed aggression (12, 13).

In general, the expression of pup-directed aggression and caregiving is mutually exclusive: male mice that engage in caregiving behaviors during a given test session do not exhibit pup-directed aggression, and vice versa (2). This dichotomy suggests that neural circuits promoting caregiving actively suppress those driving pup-directed aggression. A recent study in female mice demonstrated that *Esr1*+ neurons in the BSTpr regulate pup-directed aggression and form a reciprocal mutually inhibitory circuit with MPOA Esr1 neurons (12). However, the BSTpr contains diverse cell types (14, 15), and the specific neuronal subtype that mediates pup-directed aggression in male mice remains undetermined. Consequently, whether parental behavior-promoting circuits inhibit BSTpr neurons that drive infanticide in males remains unknown.

Our recent work demonstrated that oxytocin and vasopressin (AVP) neurons in the paraventricular hypothalamus (PVH) promote paternal caregiving behaviors in male mice. Additionally, *oxytocin receptor* (*Otr*)+ neurons in the anterior commissure nucleus (ACN) and medial preoptic nucleus function as downstream nodes that facilitate paternal behavior (16, 17). Because ACN *Otr*+ neurons are predominantly GABAergic, they may transmit inhibitory control onto infanticidal neurons.

To explore this possibility, the present study focuses on *Cartpt*+ neurons in the BSTpr. This focus was motivated by recent findings that *Cartpt*+ neurons in the medial amygdala mediate pup-directed aggression in male mice (18). Moreover, CART-peptide+ neurons in the premammillary nucleus have been implicated in maternal aggression (19), and *Cartpt* mRNA is expressed in the BSTpr (13). Together, these observations raise the possibility that BSTpr *Cartpt*+ neurons contribute to pup-directed aggression in males. Here, we show that these neurons mediate pup-directed aggression and receive direct inhibitory input from ACN neurons, and that this inhibitory connection is potentiated in father mice.

## Results

### BSTpr *Cartpt*+ neurons are activated in virgin males exposed to pups

To verify the presence of *Cartpt*+ neurons in the BSTpr and determine their neurotransmitter identities, we performed *in situ* hybridization (ISH) to detect *Cartpt* mRNA together with *vesicular glutamate transporter type II* (*vGluT2*) or *vesicular GABA transporter* (*vGAT*) (Fig. 1A). *Cartpt* mRNA was readily detected in the BSTpr. Moreover, the number of *Cartpt*+ neurons was slightly but significantly lower in fathers than in virgin males (Fig. 1B). In both groups, most *Cartpt*+ neurons co-expressed *vGAT*, indicating that these neurons are inhibitory (Fig. 1C). The proportions of *Cartpt*+ neurons co-expressing *vGluT2* or *vGAT* were comparable between fathers and virgin males (*vGluT2*, p = 0.30; *vGAT*, p = 0.83, two-tailed Mann–Whitney *U*-test).

**Fig. 1.**
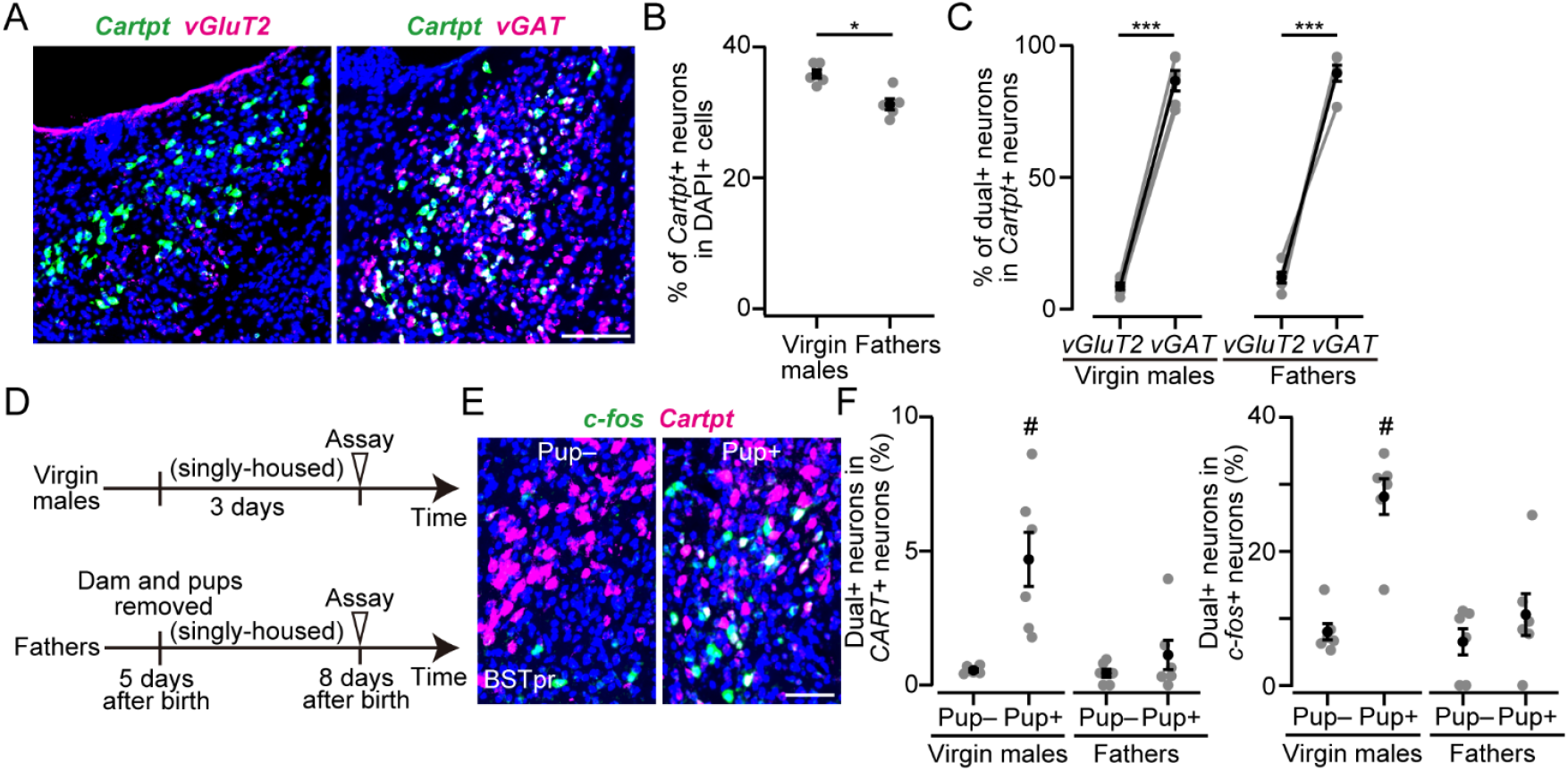
BSTpr *Cartpt*+ neurons are inhibitory neurons and are activated by pup exposure in virgin males. (A) Representative coronal sections from the father mice showing *Cartpt* (green) and *vGluT2* or *vGAT* (magenta) mRNA *in situ* staining. Blue, DAPI. Scale bar, 50 μm. (B) Fraction of DAPI+ cells expressing *Cartpt* (*Cartpt+*) is significantly smaller in father mice (*p < 0.05, two-tailed Mann–Whitney *U*-test. n = 5 each). (C) Quantification of *Cartpt+* neurons co-expressing *vGluT2* or *vGAT* (n = 5 each. ***p < 0.001, two-tailed paired *t*-test). (D) Schematic of the experiment. (E) Representative coronal sections from the virgin male mice showing *c-fos* (green) and *Cartpt* (magenta) mRNA *in situ* staining. Blue, DAPI. Scale bar, 30 μm. (F) Quantification of the fraction of *c-fos+* cells among *Cartpt+* neurons (left) and the fraction of *Cartpt+* neurons among *c-fos+* cells (right) (n = 6 each. #p < 0.01, one-way ANOVA with post-hoc Tukey HSD). Error bars, SEM.

To characterize the relationship between *Cartpt*+ GABAergic neurons and *Esr1*+ neurons in the posterior BST, we reanalyzed previously published single-nucleus RNA-sequencing datasets of *Esr1*+ neurons (14). When *Esr1+* neurons in the posterior BST were classified into 35 transcriptomic subtypes, *Cartpt* expression was enriched in four clusters (S1 Fig). One of these clusters (#18) co-expressed *Tachykinin precursor 1* (*Tac1*), which is a marker of BST neurons that trigger male sexual behavior (20), whereas the remaining three clusters did not overlap with *Tac1* expression. Together, these transcriptomic analyses indicate that BSTpr *Cartpt*+ neurons constitute a molecularly distinct subset of *Esr1*+ neurons.

The BSTpr is activated by mating and ejaculation and is modestly activated in infanticidal males (13). To examine whether BSTpr *Cartpt*+ neurons are activated when males encounter pups, we exposed animals to physically enclosed pups (see Materials and methods) and quantified *c-fos* mRNA expression as a marker of neuronal activation (Fig. 1D). Pup exposure increased the proportion of *Cartpt*+ neurons expressing *c-fos* in virgin males (Fig. 1E and 1F). This activation occurred despite the physical barrier preventing infanticidal action, which suggests that sensory exposure alone is sufficient to recruit these neurons.In contrast, fathers did not show a comparable increase in *c-fos* expression in *Cartpt*+ neurons (Fig. 1E and 1F). Collectively, these results suggest that pup exposure selectively recruits a subpopulation of BSTpr *Cartpt*+ neurons in virgin males, whereas this activation is absent in fathers.

### BSTpr *Cartpt*+ neurons promote pup-directed aggression in male mice

To determine whether BSTpr *Cartpt*+ neurons contribute to pup-directed aggression, we expressed hM3Dq, a chemogenetic activator channel, in these neurons using a previously generated *Cartpt-Cre* mouse line (21) (Fig. 2A and 2B). Clozapine-N-oxide (CNO) was administered intraperitoneally, with saline serving as a control. Because pup-directed experience can alter subsequent behavioral responses, each mouse was tested only once. The number of *Cartpt*+ neurons expressing hM3Dq did not differ significantly among groups (Fig. 2C). Consistent with previous studies (16, 17), saline-treated virgin males displayed pup-directed aggression, whereas saline-treated fathers retrieved pups (Fig. 2D). In virgin males, CNO administration modestly increased the likelihood of pup-directed aggression relative to saline controls (Fig. 2D–2G). In father mice, CNO treatment shifted behavior from caregiving to pup-directed aggression (Fig. 2D–2G). These results indicate that activation of BSTpr *Cartpt*+ neurons is sufficient to drive pup-directed aggression in males.

**Fig. 2.**
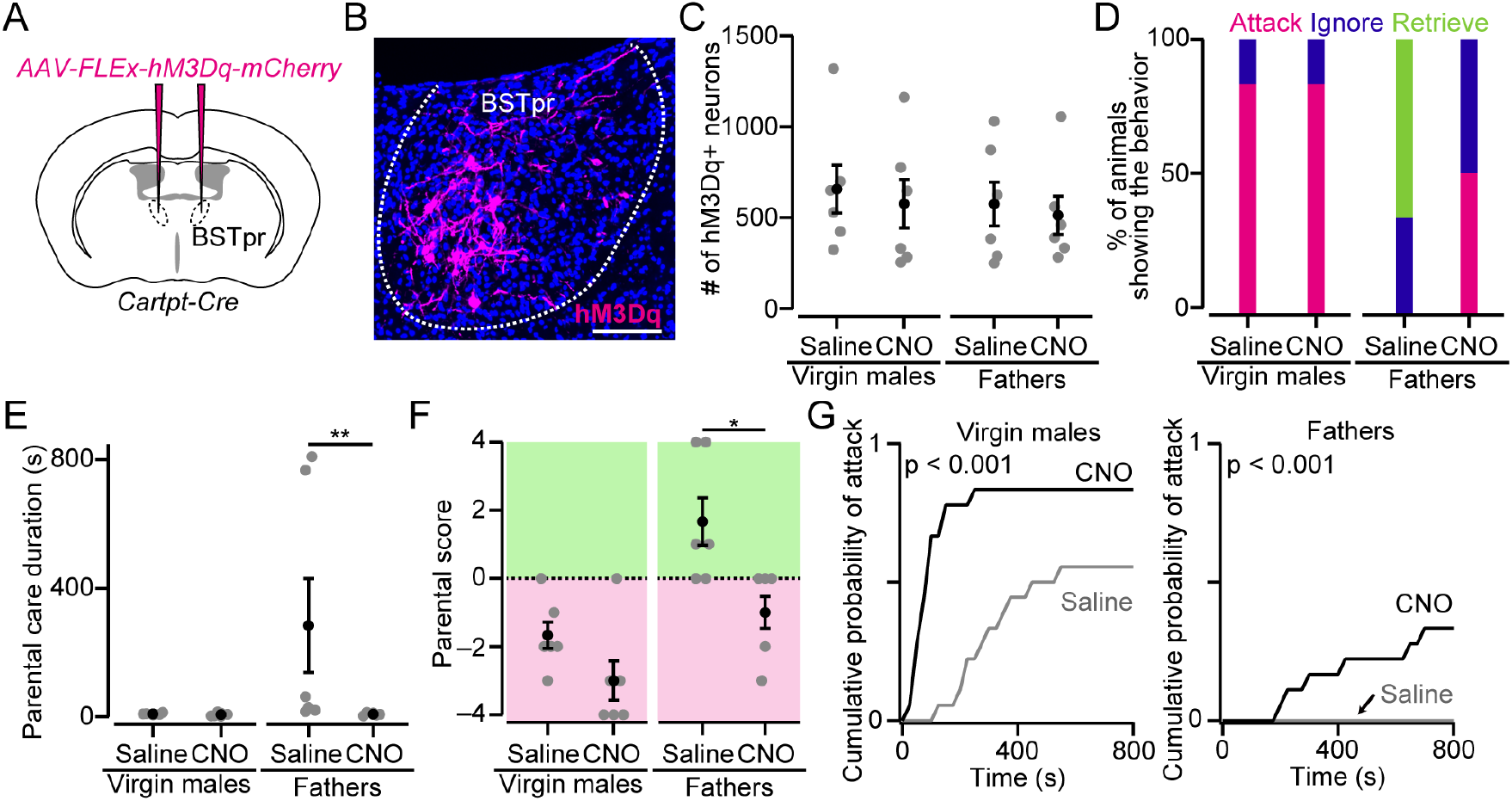
Activation of BSTpr *Cartpt*+ neurons facilitates pup-directed aggression in male mice. (A) Schematic of the virus injection. AAV *FLEx-hM3Dq-mCherry* was injected into the bilateral BSTpr of *Cartpt-Cre* male mice. (B) Representative coronal section showing the expression of hM3Dq-mCherry (magenta). Blue, DAPI. Scale bar, 50 μm. (C) The number of neurons expressing hM3Dq was not significantly different (n = 6 mice each. p > 0.57, two-tailed Mann–Whitney *U*-test). (D) Percentages of animals showing attack, ignore, or retrieve (n = 6 mice each). (E) Duration of parental care (**p < 0.01, two-tailed Mann–Whitney *U*-test). (F) Parental score (*p < 0.05, two-tailed Mann–Whitney *U*-test). (G) Cumulative probability of attack toward pups. The p value is shown in the panel (Kolmogorov–Smirnov test). Error bars, SEM.

If BSTpr *Cartpt*+ neurons are necessary for pup-directed aggression, their reduction should attenuate this behavior. To test this, we selectively ablated BSTpr *Cartpt*+ neurons by bilaterally injecting a Cre-dependent AAV encoding taCasp3-TEVp (22) into the BSTpr of *Cartpt-Cre* mice (Fig. 3A). ISH confirmed a significant reduction in *Cartpt*+ neurons following taCasp3 expression (Fig. 3B and 3C). Virgin males with reduced BSTpr *Cartpt*+ neuron numbers showed diminished pup-directed aggression (Fig. 3D–3G), and one animal displayed pup retrieval behavior, consistent with observations that suppression of aggression-associated circuits can facilitate parental behaviors (2).

**Fig. 3.**
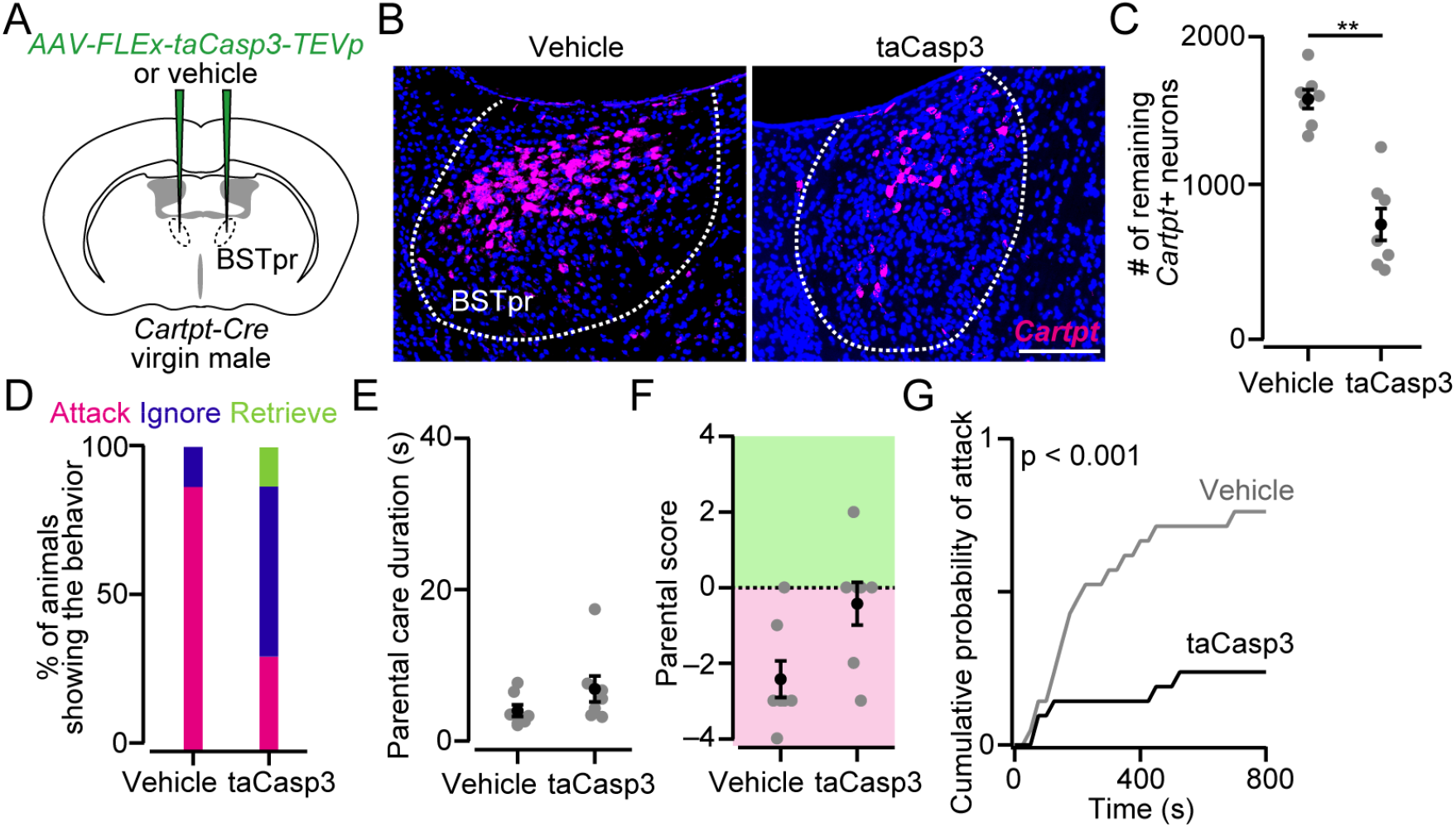
Reduction in the number of *Cartpt*+ neurons in the BSTpr suppresses pup-directed aggression in virgin males. (A) Schematic of the virus injection. AAV *FLEx-taCasp3* or vehicle (saline) was injected into the bilateral BSTpr of *Cartpt-Cre* virgin male mice. (B) Representative coronal section showing the *Cartpt* mRNA (magenta) from the mouse without (left) or with (right) taCasp3 expression. Blue, DAPI. Scale bar, 50 μm. (C) The number of neurons expressing *Cartpt* mRNA (n = 7 mice each. **p < 0.01, two-tailed Mann–Whitney *U*-test). (D) Percentages of animals showing attack, ignore, or retrieve (n = 7 mice each). (E) Duration of parental care. (F) Parental score. (G) Cumulative probability of attack toward pups. The p value is shown in the panel (Kolmogorov–Smirnov test). Error bars, SEM.

Collectively, these data demonstrate that BSTpr *Cartpt*+ neurons mediate pup-directed aggression in male mice.

### Paternal ACN provides stronger inhibitory input to BSTpr *Cartpt+* neurons

We next examined whether ACN neurons, which are predominantly GABAergic and facilitate paternal caregiving (17), provided functional synaptic input to BSTpr *Cartpt*+ neurons. To address this, we performed channelrhodopsin-2 (ChR2)-assisted circuit mapping (23). ChR2 was expressed in ACN neurons projecting to the BSTpr, and whole-cell recordings were obtained from mCherry-labeled BSTpr *Cartpt*+ neurons (Fig. 4A and 4B). Most ChR2-expressing neurons were localized within the ACN (Fig. 4C). Blue-light stimulation evoked excitatory and inhibitory postsynaptic currents in BSTpr *Cartpt*+ neurons (Fig. 4D and 4E; S2 Fig). Notably, inhibitory responses were significantly larger in fathers than in virgin males, whereas excitatory responses did not differ between groups (Fig. 4D and 4E). These findings indicate that paternal ACN neurons provide a strengthened inhibitory drive onto BSTpr *Cartpt*+ neurons, which may contribute to the suppression of pup-directed aggression in fathers.

**Fig. 4.**
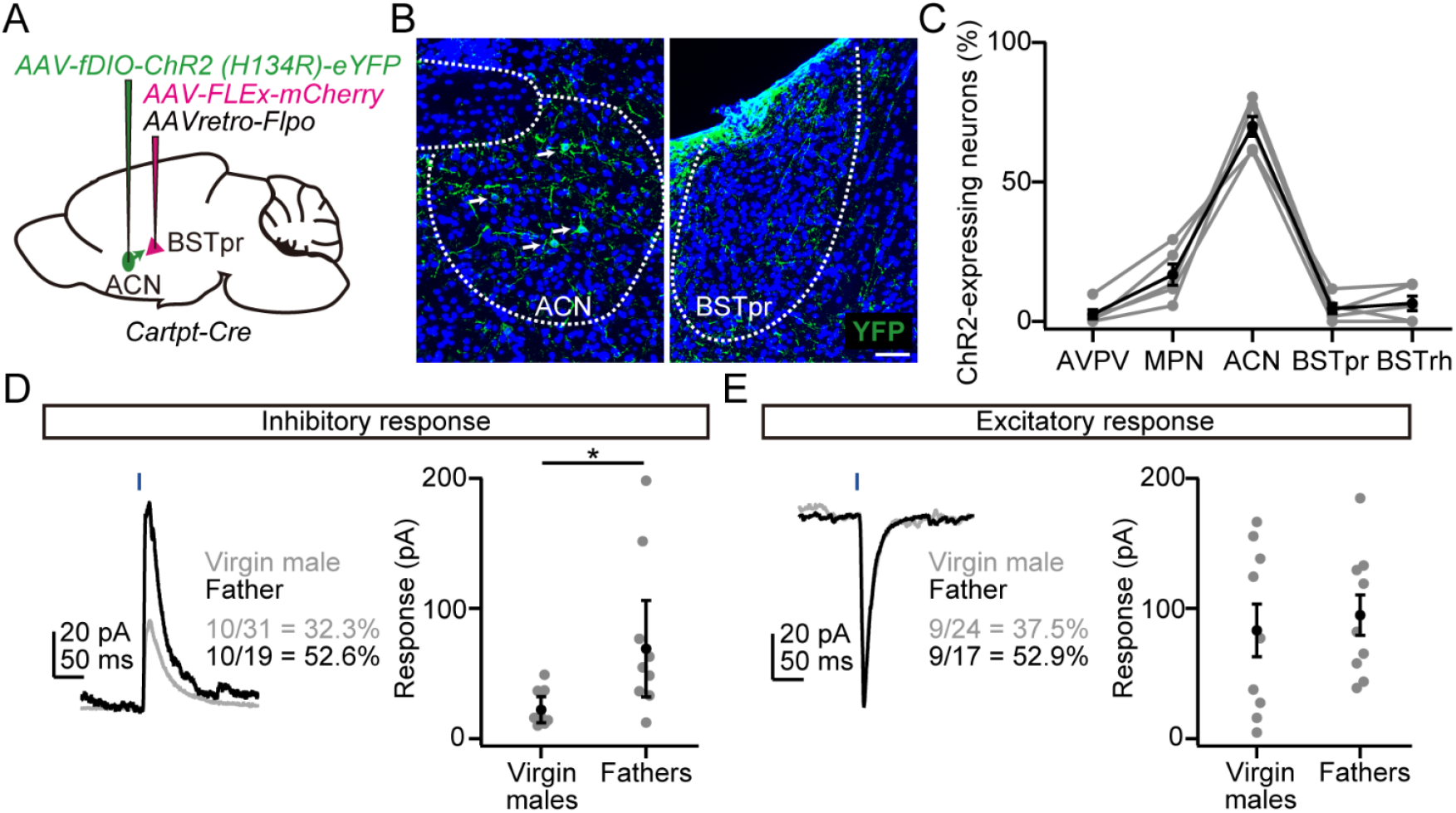
BSTpr *Cartpt*+ neurons in fathers receive stronger inhibitory inputs from ACN. (A) Schematic of the virus injection. BSTpr-projecting ACN neurons expressed ChR2, while BSTpr *Cartpt*+ neurons were labeled with mCherry. (B) Representative coronal brain sections showing the expression of ChR2-YFP (green) in the ACN (left) and BSTpr (right). Note that cell bodies expressing ChR2 (several of which are indicated by arrows) were predominantly observed in the ACN but were rare in the BSTpr. Green structure shown in BSTpr represents axonal processes. Blue, DAPI. Scale bar, 30 μm. (C) Cells expressing ChR2 were counted in the targeted nucleus (ACN) and four adjacent nuclei because our virus injection scheme may express ChR2 other than the targeted area. n = 5 virgin males. AVPV, anteroventral periventricular nucleus, MPN, medial preoptic nucleus. See main text for other abbreviations. (D, E) Representative responses from the BSTpr *Cartpt*+ neurons to the optogenetic activation of axons originating from the ACN (left). Each trace is an average of ten trials. Blue bar, light stimulation (1 ms, 3.6 mW). Percentages in each panel indicate the connected ratio. Right, the response amplitude. The inhibitory response was significantly larger in father mice than in virgin males (*p < 0.05, two-tailed Mann–Whitney *U*-test. D, n = 10 cells from eight virgin males, and n = 10 cells from five fathers; E, n = 9 cells from six virgin males, n = 9 cells from five fathers). Error bars, SEM.

## Discussion

CART peptides and *Cartpt* mRNA are distributed across multiple brain regions (24, 25), and prior studies have demonstrated *Cartpt*+ neurons in diverse physiological functions, including feeding, metabolism, nociception, and endocrine regulation (24-27). We identified a role for BSTpr *Cartpt*+ neurons in regulating pup-directed aggression in male mice. Here, we discuss the implications of these findings and the limitations of the present study.

We demonstrated that activation of BSTpr *Cartpt*+ neurons promoted pup-directed aggression and that reducing their number suppresses this behavior. Coupled with the observation that *Cartpt*+ cell numbers are reduced in fathers, these findings suggest that CART signaling contributes to the expression of pup-directed aggression in males. Despite extensive efforts since the discovery of CART, the cognate receptor remains unidentified; although a G protein-coupled receptor has been hypothesized, no receptor fully accounting for the functional diversity of CART has been confirmed (25). Identifying the receptor(s) and projection targets of BSTpr *Cartpt*+ neurons will be essential for determining how these neurons influence downstream aggression-related circuits.

The reduced *Cartpt* expression observed in fathers suggests that its expression in the BSTpr is dynamically regulated upon fatherhood. CART expression is sensitive to physiological context; for example, leptin increases CART expression in the arcuate nucleus but not consistently in the lateral hypothalamus (28, 29). In the study with prairie voles, father voles exhibit reduced body weight and body-fat composition (30), which may lower circulating leptin levels; however, such hormonal changes do not readily explain the direction observed here. Therefore, additional hormonal or transcriptional mechanisms accompanying the transition to fatherhood may modulate *Cartpt* expression in the BSTpr. Future work is required to determine the molecular pathways that downregulate *Cartpt* expression in fathers and evaluate whether this reduction is necessary for, or merely correlated with, the suppression of pup-directed aggression.

The BSTpr contains diverse cell types. The present work establishes *Cartpt-Cre* mice as a valuable tool for dissecting BSTpr neuronal subtypes responsible for pup-directed aggression. However, *Cartpt+* neurons accounted for only approximately 30% of *c-fos*+ neurons in virgin males exposed to pups (Fig. 1), suggesting that *Cartpt*-negative (*−*) neurons also contribute to pup-directed aggression. A recent study demonstrated that inhibitory *Esr1*+ neurons in the BSTpr regulate pup-directed aggression in female mice (12). Our transcriptomic reanalysis indicates that *Cartpt*+ neurons constitute a subpopulation of *Esr1*+ neurons, although the presence of *Esr1− Cartpt*+ neurons within or adjacent to the BSTpr cannot be excluded. Moreover, pup exposure activated only approximately 5% of *Cartpt+* neurons in virgin males exposed to pups (Fig. 1), suggesting that pup stimuli engage a subset of BSTpr *Cartpt+* neurons. These observations highlight the genetic and functional heterogeneity of BSTpr neurons, and underscore the need for future studies to more comprehensively characterize the molecular identities and circuit functions of BSTpr neuronal subtypes.

Upon pup exposure, oxytocin+ and AVP+ neurons in the PVH are selectively activated in father mice, which in turn activate downstream *Otr*+ neurons in the ACN (17). Because BSTpr *Cartpt*+ neurons receive stronger inhibitory input from the ACN in fathers (Fig. 4), pup exposure may suppress paternal BSTpr *Cartpt*+ neuronal activity via this inhibitory pathway. This circuit organization suggests a mechanism through which the parental state selectively inhibits aggression-promoting neurons to favor caregiving, paralleling the inhibitory interaction between MPOA *Esr1*+ neurons and BSTpr *Esr1*+ neurons described in females (12). Comparing the activity dynamics of PVH oxytocin neurons, PVH AVP neurons, ACN *Otr*+ neurons, and BSTpr *Cartpt*+ neurons during pup exposure would be a valuable next step to define the hierarchical structure and temporal logic of circuits governing pup-directed aggression and parenting.

## Materials and methods

### Ethics statements

All procedures detailed in the Materials and methods section were approved by the Institutional Animal Care and Use Committee of the RIKEN Kobe branch.

### Animals

Animals were housed under a standard 12-h light/12-h dark cycle with *ad libitum* access to food and water. Wild-type C57BL/6J mice were purchased from Japan SLC. The *Cartpt-Cre* line (Accession No. CDB0267E; listed in https://large.riken.jp/distribution/mutant-list.html) was generated previously (21). We focused exclusively on males in this study because our goal was to characterize pup-directed aggression and paternal caregiving behaviors specifically in male mice.

### Stereotactic injection

To target AAVs or vehicle (saline) into a specific brain region, we defined stereotactic coordinates for each brain region based on the Allen Mouse Brain Atlas (31). In this manuscript, the term *vehicle* refers specifically to the control solution used for microinjection into brain regions, to distinguish it from *saline*, which was used as the control for intraperitoneal CNO administration. The following coordinates were used (in mm from the bregma for anteroposterior [AP], mediolateral [ML], and dorsoventral [DV]): ACN, AP 0.0, ML 1.0, DV 4.5; BSTpr, AP −0.4, ML 0.5, DV 3.5. Mice were anesthetized with 65 mg/kg ketamine (Daiichi Sankyo) and 13 mg/kg xylazine (X1251; Sigma-Aldrich) via intraperitoneal injection and head-fixed to stereotactic equipment (Narishige). The injection volume of viruses was 200 nL at a speed of 50 nL/min, except for injections in Fig. 4, where the volume was reduced to 80 nL and the speed was 30 nL/min. After the viral or vehicle injection, the animals were returned to their home cages.

### Viral preparations

We obtained the following AAV vectors from Addgene (titer is shown as genome particles [gp] per mL): AAV serotype 8 *hSyn-FLEx-hM3Dq-mCherry* (#44361, 3.2 × 10^13^ gp/mL), AAV serotype 8 *hSyn-FLEx-mCherry* (#50459, 2.3 × 10^13^ gp/mL), and AAV retrograde *EF1a-FLPo* (#55637, 2.2 × 10^13^ gp/mL). AAV serotype 2 *EF1a-FLEx-taCasp3-TEVp* (3.4 × 10^12^ gp/mL) and AAV serotype 5 *Ef1a-fDIO-ChR2(H134R)-eYFP* (1.0 × 10^13^ gp/mL) were purchased from the University of North Carolina viral core and the Canadian Neurophotonics Platform viral vector core facility, respectively.

### Parental behavior assay

#### Assay for virgin males

The parental behavior assay was conducted as previously described (16). Briefly, 10–12-week-old male mice were housed individually for 5–7 days before the assay. Each cage contained shredded paper on wood chips. The mice build their nests with the papers, typically at a corner of the cage. Each male was tested only once. Three 5–7-day-old C57BL/6J pups that had not been exposed to any adult males before the experiment were placed in different corners where the testing male had not built its nest. The introduction of pups marked the beginning of the assay. Males were allowed to interact with the pups freely for 15 min. However, if any sign of pup-directed aggression was observed, the targeted pup was immediately rescued from the cage. In Figs. 2 and 3, experiments were conducted 3 weeks after the AAV injection. Thirty minutes before the assay, CNO (4936, Tocris) dissolved in saline was intraperitoneally injected to achieve a dose of 1 mg/kg. A saline-only injection was used as a control. The behavior of the animals was categorized into “Attack,” “Ignore,” or “Retrieve” based on the criteria previously described (16). The duration of animals undergoing grooming, crouching, or retrieving was scored as parental care duration. The parental score was calculated based on a definition described previously (17). Briefly, first, each mouse was assigned a score of 0. The number of pups retrieved or attacked by the male was added to or subtracted from this score. If the male retrieved or attacked all three pups within 2 min, one point was further added or subtracted, respectively. Consequently, the maximum value of the parental score is 4, and the minimum score is −4.

#### Assay for fathers

The behavioral assay with fathers was conducted using the same procedure as that for virgin males, with several modifications. An individually housed virgin male (10–12 weeks old) was paired with a female. The next day, the vaginal plug was checked, and only the male-female pairs that successfully formed a plug were used for further experiments. Males were allowed to cohabit with the mated female and pups until 5 days after the birth of the pups. The behavioral assay for the fathers was conducted 5 days after the birth of the pups. Mothers and pups were removed from the home cage 4–6 h before the assay, leaving only the fathers. Unfamiliar pups (pups unrelated to the resident father), prepared in the same manner as those in the assay for the virgin males, were used.

#### Assay for c-fos expression

The behavioral assay for visualizing *c-fos* expression was conducted with virgin males and fathers (Fig. 1). The assay was performed in the dark. For virgin males, each male was housed individually for 3 days before the testing. Each cage contained a metallic tea strainer. On the test day, three pups were packed into the tea strainer of the “Pup+” group, but not into the cage of the “Pup−” group. In this condition, males of the Pup+ group were able to sense the pups but could not physically contact them. This experimental paradigm was used because the assay was conducted in the dark, making it difficult for the experimenter to detect an attempted pup-directed aggression by the male mouse. Fathers were allowed to cohabit with a mated female and spend 5 days with the dam and pups. Five days after the birth of the pups, the dam and pups were removed from the home cage, and the father was left alone for 3 days before testing. Males were allowed to interact with enclosed pups for 20 min. After 20 min, males were sacrificed.

### Electrophysiology

Electrophysiological recordings were performed as previously described (16). For whole-cell patch-clamp recordings, mice were deeply anesthetized with isoflurane. Acute coronal slices (250-µm thick) were prepared using a linear slicer (Pro7N, Dosaka EM) in an ice-cold slicing solution containing (in mM): 230 sucrose, 3 KCl, 1.2 KH_2_PO_4_, 10 MgSO_4_, 10 D-glucose, 26 NaHCO_3_, 2 CaCl_2_, 5 Na ascorbate, and 3 Na pyruvate, bubbled with 95% O_2_/5% CO_2_. Slices were maintained at room temperature for at least 30 min before recording in the solution containing (mM): 92 NaCl, 20 HEPES, 2.5 KCl, 30 NaHCO_3_, 1.2 NaH_2_PO_4_, 2 CaCl_2_, 2 MgSO_4_, 25 D-glucose, 5 Na ascorbate, and 3 Na pyruvate, bubbled with 95% O_2_/5% CO_2_ (pH ∼7.3, osmolarity adjusted to 295–305 mOsm). Each slice was transferred into a submerged experimental chamber and perfused with oxygenated artificial cerebrospinal fluid (ACSF) containing (in mM): 126 NaCl, 26 NaHCO_3_, 2.5 KCl, 1.25 NaH_2_PO_4_, 1 MgSO_4_, 12.5 D-glucose, and 2 CaCl_2_, bubbled with 95% O_2_/5% CO_2_ (pH ∼7.3, osmolarity adjusted to 295–305 mOsm). Patch pipettes with a resistance of 4–6 MΩ were fabricated from thin-wall glass capillaries (1.5 mm o.d./1.12 mm i.d., TW150F-3, World Precision Instruments). The internal patch pipette solution used for voltage-clamp recordings contained (in mM): 117 cesium methanesulfonate, 20 HEPES, 0.4 EGTA, 2.8 NaCl, 0.3 Na3GTP, 4 MgATP, 10 QX314, 0.1 spermine, and 13 biocytin hydrazide (pH ∼7.3, osmolarity adjusted to 285–295 mOsm). NBQX (0373, Tocris) or gabazine (SR95531, Tocris) was dissolved at 10 μM in the ACSF. Electrophysiological recordings were made using a Multiclamp 700B amplifier (Molecular Devices), low-pass filtered at 1 kHz, and digitized at 10 kHz. A mixture of *AAV8-hSyn-FLEx-mCherry* and *AAVretro-EF1a-FLPo* was injected into the BSTpr. Two weeks later, *AAV5-Ef1a-FLEx-hChR2(H134R)-eYFP* was further injected into the bilateral ACN. Recordings were performed 2 weeks after the injection of the AAV expressing ChR2. Optogenetic stimulation for 1 ms was achieved through a 470-nm LED light (M470L3, Thorlabs). The optical intensity of the LED light was adjusted to 3.6 mW, measured at the back aperture of the objective lens (S120VC sensor, Thorlabs). Cells were held at 0 mV or −70 mV to detect inhibitory or excitatory responses, respectively. From each cell, we obtained 10–15 traces. The response amplitude was obtained by subtracting the baseline (an average of 50 ms before the light pulse) from the average of 5 ms around the peak.

### Histochemistry

Mice were anesthetized with isoflurane and perfused with phosphate buffered saline (PBS), followed by 4% paraformaldehyde (PFA) in PBS. The brain was post-fixed with 4% PFA overnight. Coronal brain sections (20-μm thick) were made using a cryostat (Leica). Fluorescent ISH was performed as previously described (16). The primers (5′–3′) used to produce RNA probes were as follows (the first one is the forward primer; the second one, the reverse primer):

*vGluT2* (part 1), 5′-TAGCTTCCTCTGTCCGTGGT; 5′-GGGCCAAAATCCTTTGTTTT

*vGluT2* (part 2), 5′-CCACCAAATCTTACGGTGCT; 5′-GGAGCATACCCCTCCCTTTA

*vGluT2* (part 3), 5′-CTCCCCCATTCACTACCTGA; 5′-GGTCAGGAGTGGTTTGCATT

*vGAT* (part 1), 5′-GCTTCCGAAACCTTTGGTG; 5′-GTACAGGCACGCGATGAG

*vGAT* (part 2), 5′-GAAGACGGGGAGGTGGTG; 5′-ATGGCCACATCGAAGAAGAC

*c-fos* (part 1), 5′-AGCGAGCAACTGAGAAGACTG; 5′-ATCTCCTCTGGGAAGCCAAG

*c-fos* (part 2), 5′-CCAGTCAAGAGCATCAGCAA; 5′-CATTCAGACCACCTCGACAA

*Cartpt* (part 1): 5′-GGACATCTACTCTGCCGTGG; 5′-TCCGGGTTGTGATGTCATCT

*Cartpt* (part 2): 5′-GCCCTGGACATCTACTCTGC; 5′-TCCGGGTTGTGATGTCATCT

In *vGluT2* ISH, a mixture of parts 1–3 was used. In *vGAT, c-fos*, and *Cartpt* ISH, a mixture of parts 1 and 2 was used. Anti-RFP (5f8; Chromotek; 1:500) and anti-rat Cy3 (712-165-153; Jackson ImmunoResearch; 1:500) were used for primary and secondary antibodies, respectively. Fluoromount (K024; Diagnostic BioSystems) was used as the mounting medium. Brain images were acquired using an Olympus BX53 microscope equipped with a 10× (N.A. 0.4) objective lens. Signal-positive cells were counted manually using the ImageJ Cell Counter plugin. Because every second section was collected, the reported number of neurons expressing hM3Dq (Fig. 2C) or *Cartpt* (Fig. 3C) was compensated (×2) from the measured value.

### Reanalysis of snRNAseq data

snRNA-seq data from Knoedler *et al*. (GSE183093) (14) were processed in R (version 4.3) using the Seurat v5 package. Because this dataset was prefiltered in the original study and contained only neuronal populations, no additional quality-control filtering or removal of non-neuronal cells was performed. Data normalization was performed using the *SCTransform* function, regressing out mitochondrial and ribosomal gene expression. Dimensionality reduction and clustering were conducted using Seurat functions *RunPCA, RunUMAP, FindNeighbors*, and *FindClusters*, using 50 principal components. Uniform Manifold Approximation and Projection (UMAP), feature plots, and dot plots were generated after subsetting male samples. The number of clusters was retained as defined in the original publication.

### Data analysis

All mean values are reported as the mean ± standard error of the mean (SEM). The statistical details of each experiment, including the statistical tests used, the exact value of n, and what n represents, are shown in each figure legend. The p-values are shown in each figure legend or panel; nonsignificant values are not noted.

## Author contributions

Conceptualization: K.I.

Funding acquisition: K.I. and K.M.

Investigation: K.I., H.S., and M.H.

Writing – original draft: K.I. and K.M.

Writing – review and editing: K.I. and K.M.

## Acknowledgments

We thank RIKEN BDR animal facility for animal care and *in vitro* fertilization, members of the Miyamichi Laboratory for the critical reading of the manuscript, Addgene, the Canadian Neurophotonics Platform viral vector core, and the University of North Carolina Vector Core for the AAV production.

## Supporting information

**S1 Fig.**
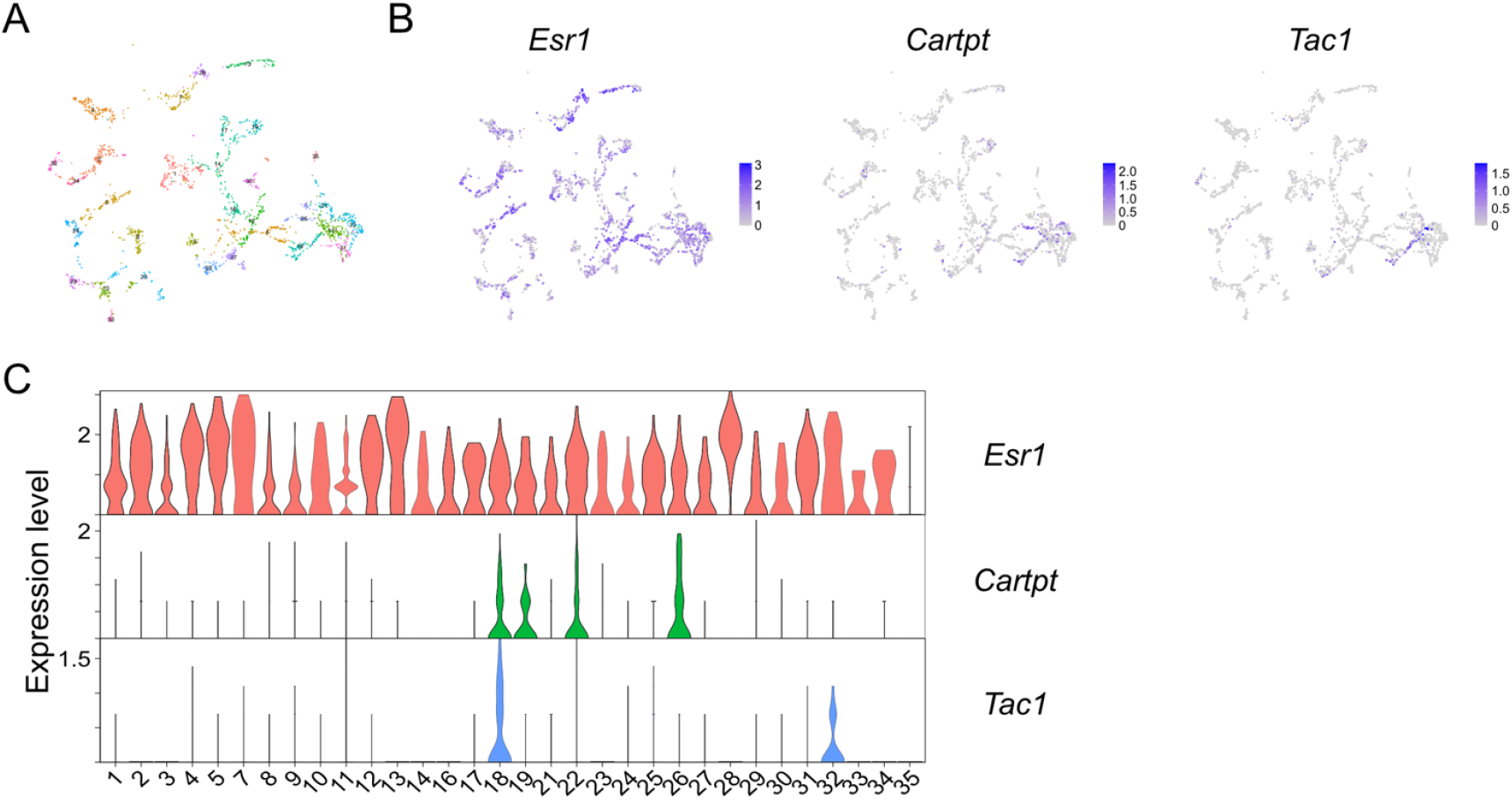
A, UMAP representation showing *Esr1*+ inhibitory neuron clusters in BST based on the reanalysis of snRNAseq dataset (14). B, Expression levels of *Esr1, Cartpt*, and *Tac1* among BST *Esr1*+ inhibitory neurons. C, Violin plots showing the expression levels of *Esr1, Cartpt*, and *Tac1* among BST *Esr1*+ inhibitory neuron clusters.

**S2 Fig.**
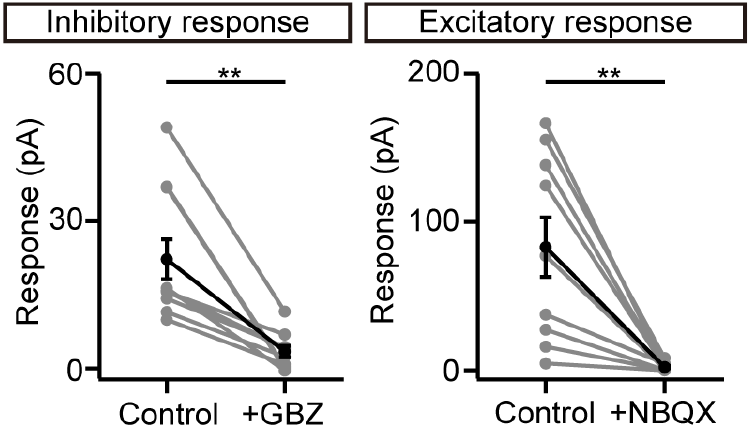
Significant decrease in the inhibitory (left) or excitatory (right) responses under optogenetic stimulation after the application of gabazine (GBZ) or NBQX (left, n = 10 cells from eight virgin males, right, n = 9 cells from six virgin males. **p < 0.01, paired *t*-test). Control data are shown in Fig. 4D and 4E. Error bars, SEM.

